# The consequences of mixed-mode transmission for disease prevalence

**DOI:** 10.64898/2026.07.22.740081

**Authors:** Irish Amundson, Janis Antonovics

## Abstract

Host-parasite relationships are defined by transmission. Transmission takes diverse forms, with horizontal transmission modes generally characterized as density- or frequency-dependent. However, many observed parasites display mixed-mode transmission using both density- and frequency-dependent routes. To investigate how mixed-mode transmission impacts epidemics, we assessed infection prevalence under single- and mixed-mode transmission when there was a linear trade-off between the probability of infection by the two modes. Our results show that the prevalence of a parasite with mixed-mode transmission can be greater than one with a single-mode only when density- and frequency-dependent routes are associated with different effects of the parasite on host fitness. This work shows that mixed transmission modes per se may not necessarily increase disease prevalence, and that substituting two transmission modes for a single one may result in higher prevalence under limited conditions.

## Introduction

Public health concern has generated new interest in ecological and evolutionary processes precipitating epidemics. Parasite transmission is a major factor predicting epidemic severity and duration. Transmission is also a primary determinant of parasite fitness, as parasites must infect new hosts to persist. In this study, we investigate the ecological consequences of parasite transmission strategies, focussing on the performance of parasites with mixed transmission modes. In population models of disease dynamics, two frameworks have been especially fruitful in furthering the study of horizontal transmission. The first and traditionally assumed assumes that transmission events increase with host density; we refer to this as density-dependent transmission [1]. The second assumes that transmission is independent of host density and instead increases with the proportion of infected hosts; we refer to this as frequency-dependent transmission [2]. Models of density-dependent transmission tend to be a good approximation for direct non-sexual contact or air-borne transmission, while frequency-dependent transmission approximates sexual and vector-borne transmission [2,3].

While density- and frequency-dependent transmission are often treated as mutually exclusive, many diseases cannot be neatly sorted into one or the other mode based on empirical observations [4–9]. Ebolavirus in humans, anther-smut in plants, foot-and-mouth disease in cattle, human herpesvirus 8, and syphilis all demonstrate a mixture of these two modes [10–15]. We typically consider sexually transmitted diseases to be frequency-dependent, but Lockhart et al. 1996 [16], found that of 108 sexually transmitted diseases of animals, 87 could also transmit by alternative means, many of which involve social contact and are thus likely density-dependent. In plants, anther-smut disease causes flowers to produce fungal spores rather than pollen. Detailed experimental studies demonstrate that flowers spores to nearby plants in a density-dependent fashion, while pollinators carry spores from one plant to another in a frequency-dependent fashion [11,17,18]. Here, we refer to such transmission as mixed-mode, meaning that transmission can happen through multiple routes, with at least one route dependent on density and one route dependent on frequency. We contrast mixed-mode transmission with single-mode transmission, which is strictly density-dependent or frequency-dependent. Prior work suggests that mixed-mode transmission has major impacts on epidemic dynamics; if even a small degree of mixed-mode transmission occurs, a single-mode model may not accurately predict epidemics, parasite persistence, or host extinction. Mixed-mode transmission may increase the parameter space in which both endemicity and parasite-driven extinction are possible [19].

Transmission mode may also influence the fitness consequences disease infection. At a broad, comparative level, diseases that result from aerial transmission are generally density-dependent and tend to lower host survival, while sexually transmitted diseases are usually frequency-dependent and often cause host sterility [20]. This phenomenon is observed in, for example, ocular and genital chlamydia [21,22], cutaneous and mucosal HPVs [23], and diseases caused by *Treponema pallidum* (yaws and syphilis) [24]. Sexually transmitted parasites may benefit evolutionarily when they impose sterility rather than mortality costs on hosts [25,26]. When allocation to sexual transmission increases host sterility and allocation to social transmission increases host mortality, Thrall et al. 1998 argued that mixed transmission strategies could increase parasite fitness [25]. In anther-smut, frequency-dependent transmission permits some reproduction prior to infection, while density-dependent transmission to seedlings results in lifetime sterility and may increase mortality [27,28].

A fundamental question remains: do parasites with mixed-mode transmission spread more effectively than parasites with a single mode? Here, we address this question by evaluating the prevalence of a mixed-mode disease as compared to its single-mode counterparts where investment in one transmission mode “trades-off” with investment in the other. We find that the prevalence of a disease with a mixed transmission mode cannot be predicted by a simple additive relationship of the single modes. Moreover, we show that mixed-mode transmission can only result in higher prevalence than the single modes when the component modes incur distinct fitness consequences.

## Methods

We present a deterministic model [2,29–32], where hosts are either “susceptible”, or “infected.” The number of susceptible hosts, the number of infected hosts, and the total number of hosts are represented with *S, I*, and *N*, respectively. Hosts flow in and out of our model’s “compartments” through demographic processes, and transition from “susceptible” to “infected” via infection through two transmission modes. We assume there is no recovery from infection.

Demographics in our models include birth and death. Susceptible hosts are born at rate *b* and die at rate *d*. Throughout, we assume that the birthrate exceeds the death rate: *b* > *d*. The total number of hosts in the environment is limited to a carry-capacity *k*. When the population reaches *k*, no new births occur, i.e. we assume that *b*(1 − *N*/*k*) cannot be less than zero. See Table 1 for a complete list of parameters, their range of values, and other assumptions.

**Table 1.** All parameter definitions alongside bounds, assumptions, and examples of usage in the parameter space explored.

| Parameter | Definition | Bounds |  |
| --- | --- | --- | --- |
| $b$ | birthrate | $0 \leq b \leq 2$ | When $b = 1$, each host produces one additional susceptible host. |
| $d$ | deathrate | $0 \leq d \leq 1$ | When $d = 0$, no hosts die. When $d = 1$, all hosts die. |
| $k$ | carrying-capacity | $k = 200$,<br>$N/k \leq 1$ | When $N/k = 0$, $bS$ births occur. When $N/k \geq 1$, no births occur. |
| $\beta_f$ | frequency-dependent transmission coefficient | $0.01 \leq \beta_f \leq 0.9$ | In one simulation, when $\beta_f = 0.6$, prevalence was 0.5. |
| $\beta_d$ | density-dependent transmission coefficient | $0.0001 \leq \beta_d \leq 0.1$ | In one simulation, when $\beta_d = 0.06$, prevalence was 0.5. |
| $c$ | relative investment in frequency-dependent transmission | $0 \leq c \leq 1$ | When $c = 1$, transmission is strictly frequency-dependent. When $c = 0$, transmission is strictly density-dependent. |

**Table 2.**
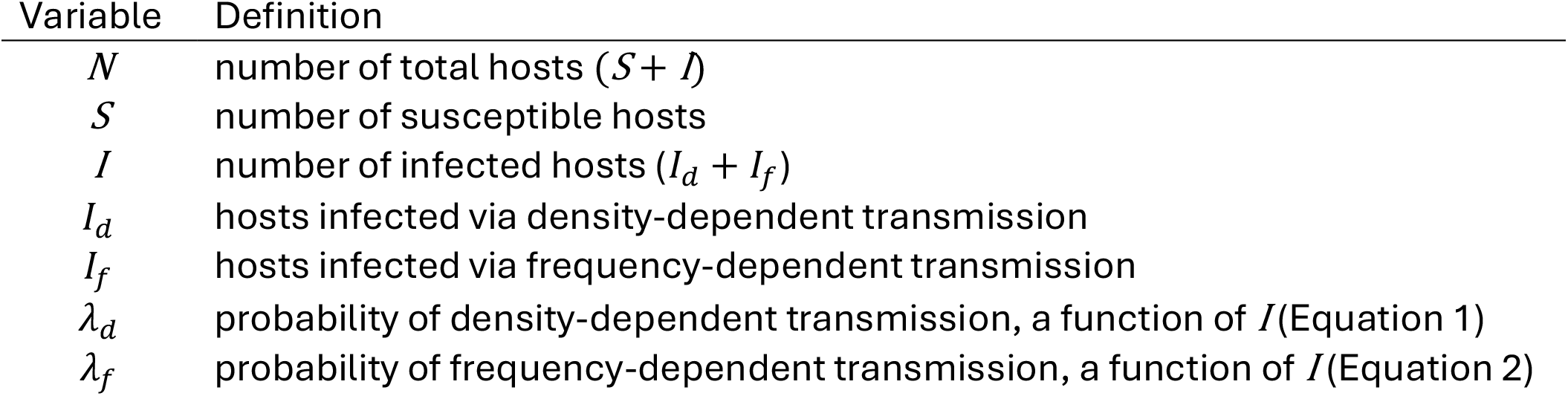
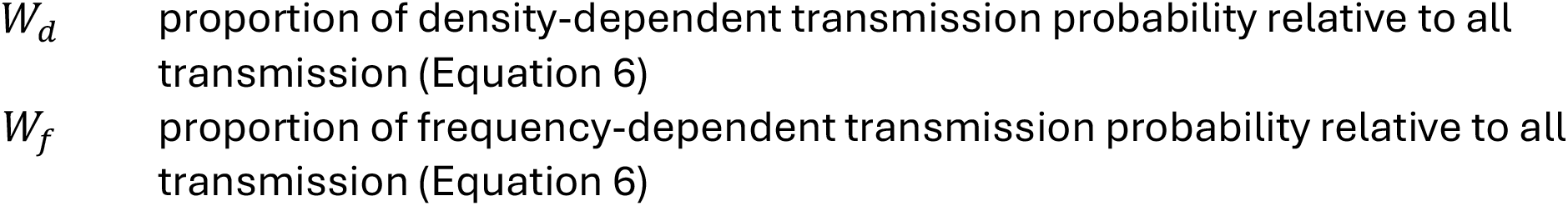
All variable definitions. Variable Definition.

Hosts transition from the “susceptible” compartment to the “infected” compartment according to the rate of transmission. Transmission occurs when a susceptible host has contact with an infected host and that contact results in infection. Because we are concerned with two modes of contact with infected hosts (density- and frequency-dependent), our models include two different transmission terms. Under density-dependent transmission, hosts contact one another randomly in space. The contact rate between susceptible and infected hosts is thus proportional to *SI*. Only a fraction of these contacts results in infection. We accordingly multiply the contact rate by a transmission coefficient, *β*_*d*_ [1]. Thus, the rate of density-dependent transmission is *β*_*d*_*SI*. Under frequency-dependent transmission, the effect of density on contact is overcome by some active mechanism, such as a mosquito searching for a bloodmeal or a host searching for a mate. Thus, for every *S*, the probability of a contact is the proportion of infected hosts, *I*/*N*, and infection occurs at a rate *β*_*f*_. This makes the rate of frequency-dependent transmission *β*_*f*_*S I*/*N* [2].

To implement a simple trade-off in the parasite’s investment in each mode, we express allocation to frequency-dependent transmission as a proportion, *c*. The investment in density-dependent transmission is then 1 - *c*. The rate of frequency-dependent transmission is then *cβ*_*f*_*S I*/*N*, while the rate of density-dependent transmission is (1 − *c*)*β*_*d*_*SI*. We express transmission in exponential Nicholson-Bailey form [33,34], where the probability of infection is 1 − *e*^*-f(x)*^ such that *f*(*x*) gives the frequency- or density-dependent transmission rate per *S*. We use this form because it expresses each transmission mode in equivalent units and also allows us to investigate the model numerically over a large range using randomly drawn parameter values while avoiding parameter spaces in which new infections outnumber available susceptible hosts. We represent the two transmission functions using *λ*_*f*_ and *λ*_*d*_ for frequency- and density-dependent transmission, respectively.

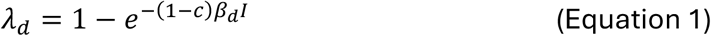

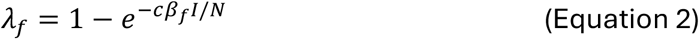

Because we express transmission as a probability, there are three possibilities for transmission: 1) a host may be infected through frequency-dependent transmission, but not density-dependent transmission, given by *λ*_*f*_*(1* − *λ*_*d*_); 2) a host may be infected through density-dependent transmission, but not frequency-dependent transmission, *λ*_*d*_*(1* − *λ*_*f*_); or 3) a host may be infected through both simultaneously, given by *λ*_*f*_*λ*_*d*_. When this happens, we assume that only one infection occurs. The type of infection results in proportion to its probability of infection over the total infection probability. This proportion is given by Equation 3, where *i* is either *f* or *d*. The respective *W* thus assigns the dually exposed hosts proportionately to either the *I*_*f*_ or *I*_*d*_ compartment.

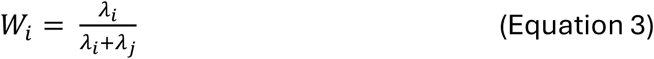

We constructed two models that differed in whether the mode of transmission affects the fitness consequences for the host. Fitness consequences of infection can take the form of increased mortality (*d*_*d*_, *d*_*f*_), decreased fertility (*b*_*d*_, *b*_*f*_), or both. Infection is never beneficial to the host such that *d*_*d*_ and *d*_*f*_ ≥ *d*, and *b*_*d*_ and *b*_*f*_ ≤ *b*. In Model A, infected hosts experience all fitness consequences of disease regardless of the mode through which they were infected. Thus, in Model A, *d*_*d*_ *= d*_*f*_ and *b*_*d*_ *= b*_*f*_. In Model B, the mode of infection determines the fitness consequence of infection. Hosts infected through frequency-dependent transmission experience only fertility consequences; hosts infected through density-dependent transmission experience only mortality consequences. Accordingly, in Model B, *d*_*f*_ *= d* and *b*_*d*_ *= b*.

Equation 4 describes the change in the number of susceptible hosts through time. Susceptible hosts are gained through density-dependent births. They are lost through death, as well as infection.

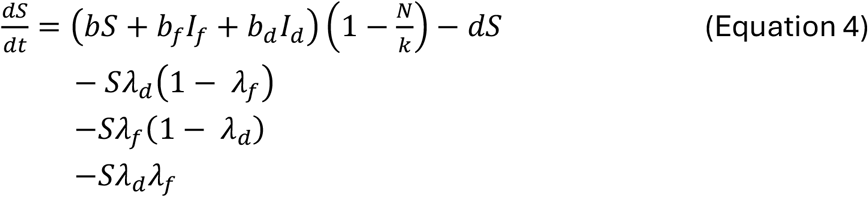

Equations 5 and 6 describe the accompanying change in the number of hosts infected through frequency- and density-dependent transmission, respectively. Infected hosts are lost through death and gained by new transmission.

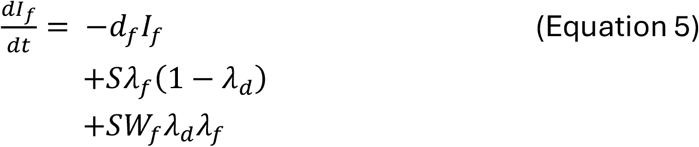

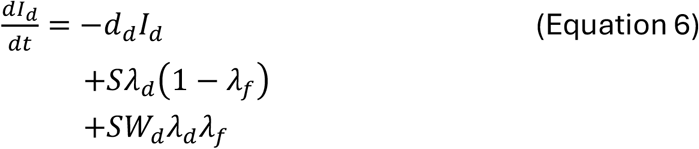

### Implementation

We ran simulation models in R version 4.5.1 and used the package “deSolve” (version 1.40) to solve ODEs numerically [35]. We analysed the outcomes of each model by varying parameters and initial values from a uniform distribution (Table 1). We avoided parameter space that overwhelmingly resulted in extinction of the host or parasite. We held the carrying capacity constant for all simulations (*k* = 200). For each parameter sampling, we varied investment, *c*, from 0 to 1 by increments of 0.05. We drew 1000 parameter sets for each model. We ran simulations for 100 timesteps after which we evaluated whether they reached equilibrium, defined as when *S* and *I* had changed by less than 10^-6^ over ten timesteps. We continued simulations until they reached equilibrium, then evaluated prevalence, the proportion of infected hosts in the population 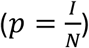, as well as the contribution of each transmission mode to prevalence in the absence of the other mode.

For the figures shown, we chose representative simulation outcomes and present them as replacement series graphs, akin to their use in showing the results of experiments where two plant species are grown both together in various proportions and in monoculture. By analogy, we use the terms “under-” and “over-prevalent” to describe results where mixed-mode disease prevalence is lower or higher than the average of the single-modes in pure culture by allocation to each mode. We use the term “super-prevalent” when the prevalence of any mixed-mode is greater than both single-modes. We compare results to an “expected” prevalence at which there is neither under-nor over-prevalence. To determine which parameters most influenced super-prevalence, we took a representative super-prevalent Model B simulation and modified the parameters from low to high.

## Results

We first asked whether a simulation’s prevalence was greater or less than expected. We then asked whether the simulation had a mixed-mode prevalence greater than the prevalence for both single-modes, i.e., super-prevalence. We compared simulation where infection by the two modes had the same (Model A) or distinct (Model B) fitness consequences. We found that Model A simulations could be over- or under-prevalent, but never super-prevalent, whereas Model B simulations were often super-prevalent.

### Model A

In Model A, fitness consequences do not vary between infected hosts, regardless of how infection was transmitted. We found that under mixed-mode transmission, prevalence is not proportional to investment and is never super-prevalent, such that maximum prevalence is always associated with either strict density- or frequency-dependence. Representative instances of Model A are shown in Figure 1. Mixed-modes could be over-prevalent (Figure 1A, 1B), or under-prevalent (Figure 1C, 1D). We observed these two results approximately equally frequently in the parameter space we examined. Frequently, a single-mode alone (Figure 1, shown by grey lines) is insufficient to spread disease especially at low allocation to that mode, and adding the alternative mode greatly increases prevalence.

**Figure 1.**
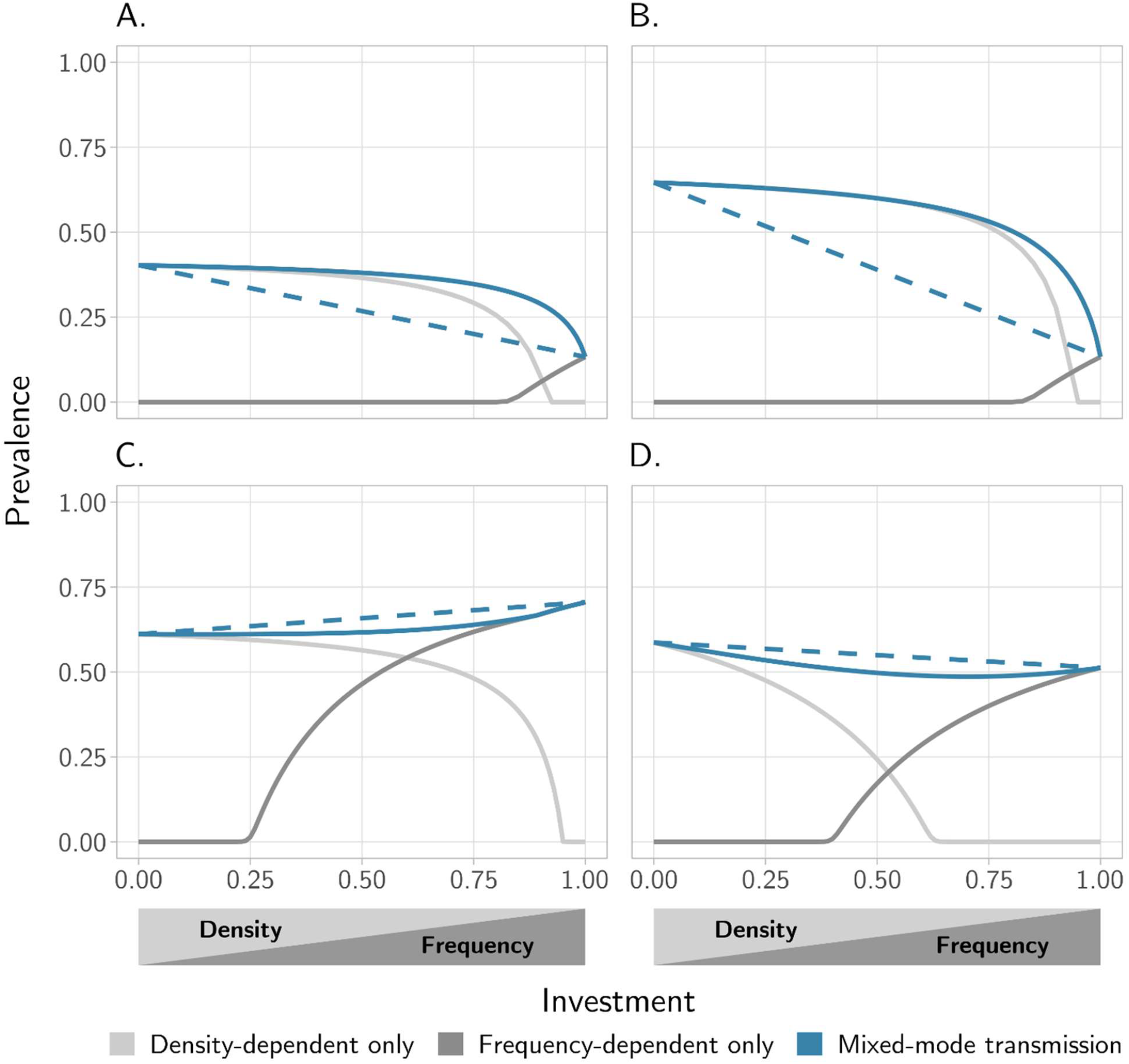
Representative instances of Model A. The x-axis is proportion of investment, *c*, in frequency-dependent transmission. Solid blue lines show prevalence of disease with mixed-mode transmission. Gray lines show prevalence for density- or frequency-dependent transmission alone. Dashed blue lines show expectation if prevalence was predicted based on values for each transmission mode separately (values at *c* = 0 and 1). For all panels, *k* = 200, *b*_*f*_ = *b*_*d*_, and *d*_*f*_ = *d*_*d*_. **A:** *b* = 0.6, *d* = 0.3, *β*_*f*_ = 0.6, *β*_*d*_ = 0.06, *d*_*d*_ = 0.5, *b*_*f*_ = 0.12. **B:** *b* = 1.0, *d* = 0.3, *β*_*f*_ = 0.6, *β*_*d*_ = 0.06, *d*_*d*_ = 0.5, *b*_*f*_ = 0.2. **C:** *b* = 0.2, *d* = 0.1, *β*_*f*_ = 0.6, *β*_*d*_ = 0.03, *d*_*d*_ = 0.15, *b*_*f*_ = 0.1. **D:** *b* = 2.0, *d* = 0.3, *β*_*f*_ = 0.8, *β*_*d*_ = 0.005, *d*_*d*_ = 0.32, *b*_*f*_ = 0.4.

### Model B

In Model B, the mode of transmission affects the fitness consequence the host experiences. In this model, mixed-mode transmission results in super-prevalence for the majority of simulations (Figure 2A, 2B), though some simulations were over-prevalent without super-prevalence (Figure 2C). Simulations were only very rarely under-prevalent (Figure 2D), and such simulations tended to have low overall prevalence. Similar parameter sets were used to generate Panels A:C for Figures 1 and 2. The qualitatively different result from the same parameter set demonstrates that super-prevalence is due to the difference in outcomes between the models. When the mode of transmission affects the fitness consequence in the manner described by Model B, super-prevalence may result.

**Figure 2.**
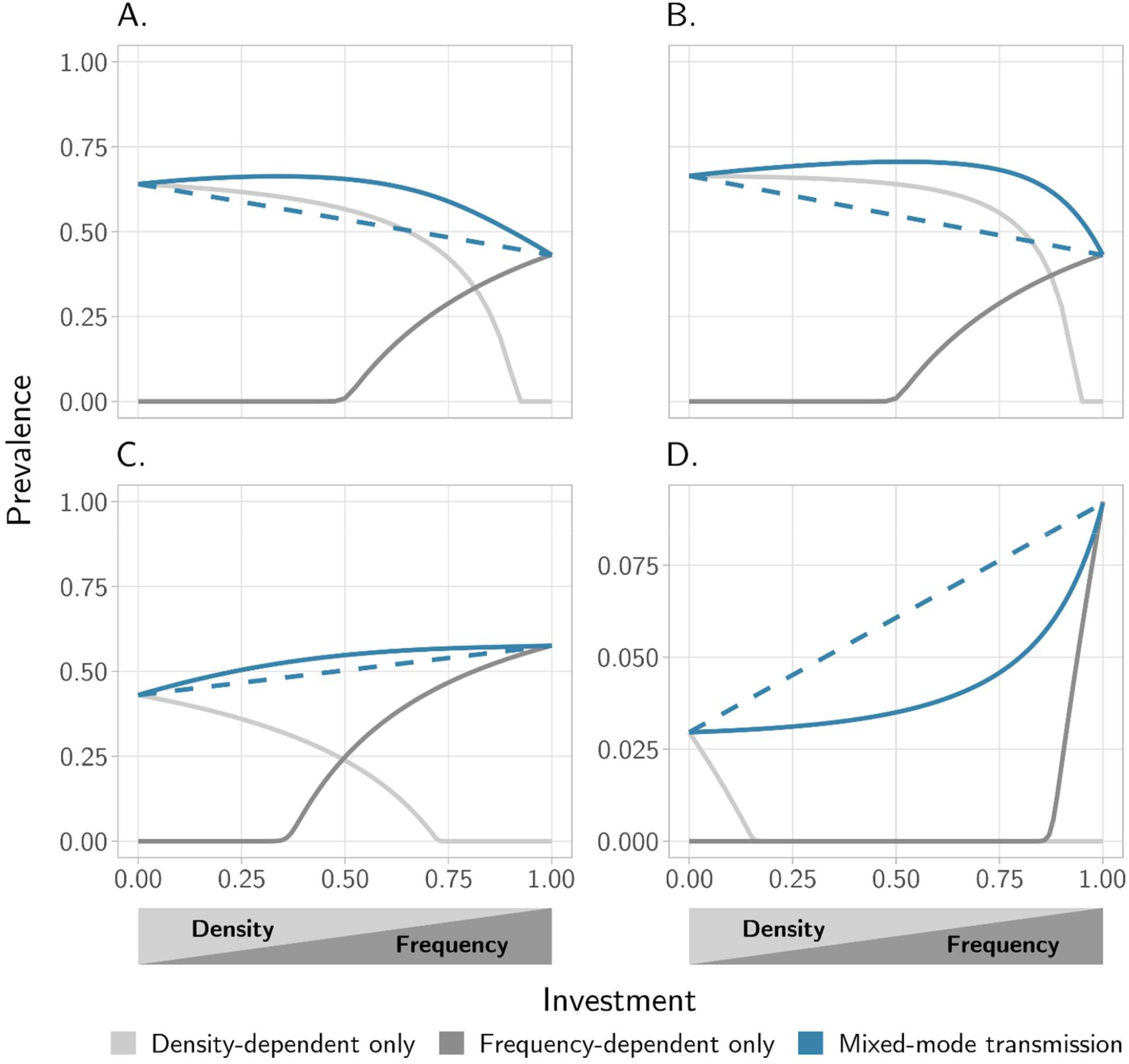
Representative instances of Model B. Notation and labelling are presented as in Figure 1. Under-prevalent results are observed in simulations with low overall prevalences and only rarely; thus, Panel D is presented at a different scale to other panels. For all panels, *k* = 200, *b*_*d*_ = *b*, and *d*_*f*_ = *d*. **A:** *b* = 0.6, *d* = 0.3, *β*_*f*_ = 0.6, *β*_*d*_ = 0.06, *d*_*d*_ = 0.5, *b*_*f*_ = 0.12. **B:** *b* = 1.0, *d* = 0.3, *β*_*f*_ = 0.6, *β*_*d*_ = 0.06, *d*_*d*_ = 0.5, *b*_*f*_ = 0.2. **C:** *b* = 0.6, *d* = 0.2, *β*_*f*_ = 0.55, *β*_*d*_ = 0.015, *d*_*d*_ = 0.55, *b*_*f*_ = 0.18. **D:** *b* = 0.75, *d* = 0.7, *β*_*f*_ = 0.8, *β*_*d*_ = 0.08, *d*_*d*_ = 0.9, *b*_*f*_ = 0.375.

#### Maximizing prevalence

To determine what parameter space tended to yield super-prevalence, we varied parameters (*b, b*_*f*_, *d*, and *d*_*d*_) iteratively using the parameter space presented in Figure 2A as a baseline. For each resulting simulation, we noted the investment, *c*, associated with maximum prevalence. This investment for each simulation is presented in a phase diagram (Figure 3). We found that super-prevalence tended to occur when birthrates were intermediate or high (Figure 3). We also found super-prevalence was more likely when infection caused only intermediate or mild sterility. Highly sterilizing parasites tended to have maximal prevalence when they invested more in density-dependent transmission, while highly deadly parasites tend to be maximally prevalent when they invested more in frequency-dependent transmission. As disease-induced mortality increased, maximally prevalent parasites were increasingly frequency-dependent; as disease-induced sterility increased, maximally prevalent parasites were increasingly density-dependent (Figure 3).

**Figure 3.**
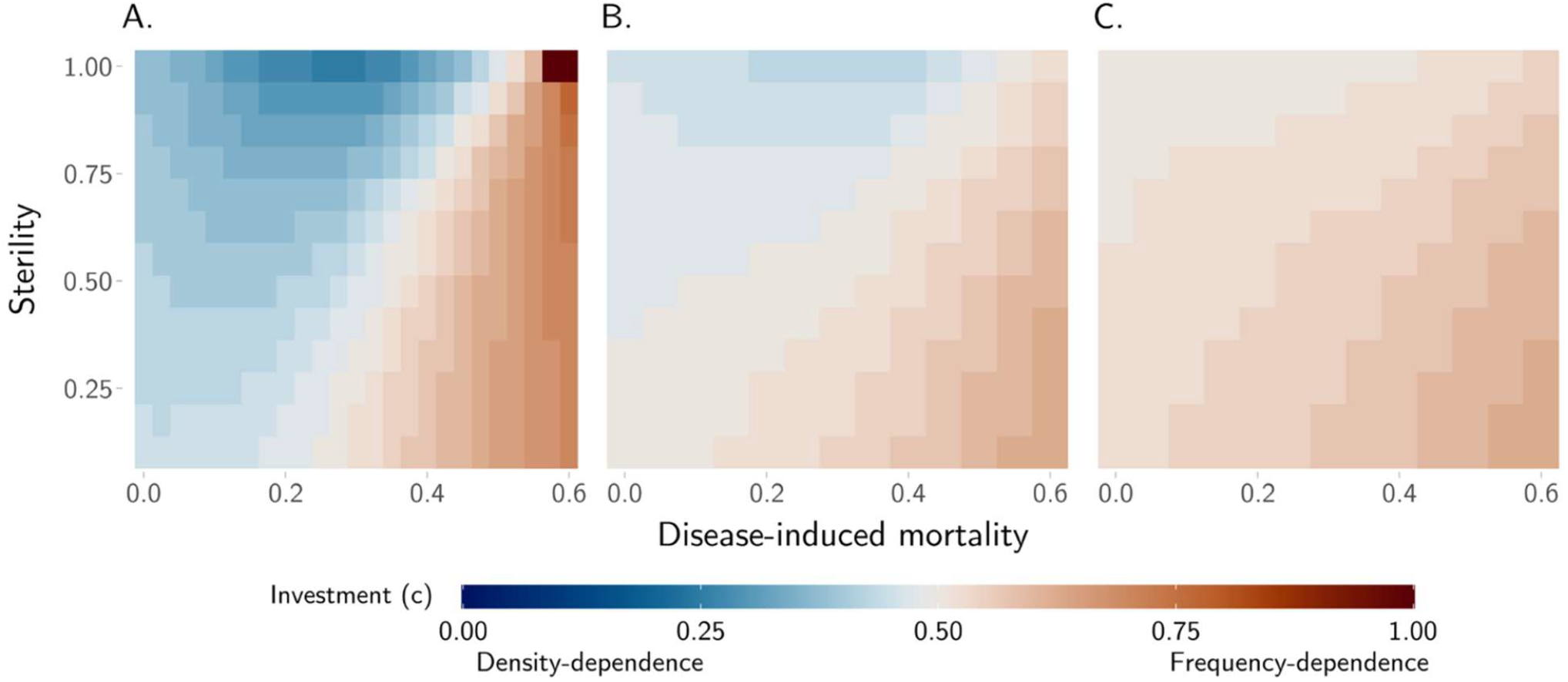
Phase diagram for Model B showing level of investment in transmission mode that achieves maximum disease prevalence when frequency-dependent transmission causes different levels of sterility and density-dependent transmission causes different levels of mortality. For a given pair of values for disease-induced mortality and sterility, the maximally prevalent disease’s investment with represented by colour. Intermediate colours show super-prevalent results. For all panels, *d* = 0.3, *β*_*f*_ = 0.6, and *β*_*d*_ = 0.06. Panels differ only by birthrate (**A:** *b* = 0.6, **B:** *b* = 0.8, **C:** *b* = 1.0).

## Discussion

Our results show that mixed-mode transmission can result in under-, over, and super-prevalence relative to single-mode transmission. This super-prevalence is present only when the two transmission routes inflict different fitness effects. Under this condition, mixed-mode transmission is the most prevalent strategy when host birthrate is moderate or high. When birthrate is low, super-prevalence is more likely when infection causes moderate or mild sterility.

In both models, mixed-mode transmission did not support a linear relationship between investment and prevalence. Rather, mixed-mode transmission caused under- or over-prevalence. Our findings suggest that, when present, mixed-mode transmission should be modelled explicitly and one should not assume a linear relationship between prevalence with only one mode and total prevalence, nor between prevalence and allocation to individual modes. Without explicit modelling, there is a risk of drawing substantively inaccurate conclusions regarding diseases that use mixed-mode transmission. Our results validate prior evidence that complexities of transmission mode need to be taken into account for correct epidemic predictions [6,36–39].

We describe investment as a linear trade-off in transmission mode. This trade-off shape is likely for natural systems in which investment is controlled by a single trait, such as the dispersal mode of the infective stages. For example, while many *Microbotryum* species produce spores in the anthers of the host plant and transmit through pollinator vectors, a fraction of the spores is likely to be wind dispersed. The pathogen *M. majus* on *Silene otites* produces spores in the whole flower and is likely primarily wind-dispersed [40]. We have personally observed that *M. majus* spores are dustier than other common anther-smuts (e.g., *M. lychnidis-dioicae*). Where multiple traits control investment, other trade-off shapes are likely. Here, we chose a linear relationship as a simple shape to address our question in an idealized, simulated system, though other shapes could be explored in future work. Importantly, this linear trade-off does not necessarily translate into a proportional relationship between density- and frequency-dependent transmission rates. The two rates are only proportional when the transmission terms are equivalent (i.e. when *β*_*f*_ *I*/*N* = *β*_*d*_*I*).

Alternatively, under- and over-prevalence may arise from the fact that population size responds differently to frequency- and density-dependent disease transmission [2,32]. Density-dependent disease is unable to spread below a certain host population size threshold. Such a disease also tends to impose a cap on host population size, which may be well below the environmental carrying capacity. Frequency-dependent disease has no such relationship with host population size. Thus, it is possible that prevalence varies in proportion to some function of both investment and the resulting effect on population size.

In addition to simple over-prevalence, we also found super-prevalence. When the type of fitness consequence varies by transmission route, parasites with mixed-mode strategies can achieve higher prevalence (Figure 2). In Model B, we consider the scenario in which frequency-dependent transmission causes sterility and density-dependent transmission causes mortality. We devised this scenario based on observation of natural disease systems [16,20,21,41]. We did not consider the reverse relationship, with frequency-dependent transmission causing mortality and density-dependent causing sterility; this is a fruitful avenue for future work. Our results here are consistent with Thrall et al. 1998 [25], in which the authors investigated models with two pathogen strains differing in whether they were transmitted sexually (frequency-dependent) or socially (density-dependent). Similar to our work, the authors found that mixed social and sexual contact could be favoured when fitness consequences varied, such that increasing sexual transmission decreased fertility and increasing social transmission increased mortality. When fitness consequences did not vary, only purely social or sexual transmission was favoured.

Our findings have potentially important evolutionary implications. Prevalence may be a reasonable proxy for parasite fitness at equilibrium. From this perspective, our work implies that mixed-mode transmission may constitute a parasite fitness peak when fitness consequences differ by transmission mode. We expect mixed-mode transmission to be favoured specifically when birthrate is moderate or high and when disease-induced sterility is mild (Figure 3). Intermediate and high birthrates may keep the frequency- and density-dependent contact rates close together, maintaining mixed strategies. The point at which those contact rates are equivalent is analogous to the “social-sexual crossover point” described in previous work as the point at which social and sexual contact occur at the same rate [20,25,41]. Selection may thus favour a mixed strategy or transmission mode polymorphism [25,41], perhaps such as that observed in closely related STD’s and nonsexual diseases [21–23,41]

In summary, our work assesses the ecological consequences of a simple model of mixed-mode transmission. We establish expectations for the behaviour of mixed-mode transmission. We find that mixed-mode transmission confers higher prevalence than single-mode strategies when transmission routes lead to different fitness consequences for the host.

## Acknowledgments

We thank Michael Hood for the suggestion to use a replacement series approach to understanding mixed-mode transmission.

We also thank Mandy Gibson, Emme Bruns, and Louis Bubrig for helpful comments on earlier drafts of this paper and discussion of these ideas.

This work was based on work supported by the NSF-GRFP to IA (Grant No. 1842490) and the NSF-NIH-USDA Ecology and Evolution of Infectious Diseases Program to JA (R01GM122061).

## References

1. May RM, Anderson RM. 1979 Population biology of infectious diseases: part I. Nature 280, 361–367. (doi:10.1038/280361a0)

2. Getz WM, Pickering J. 1983 Epidemic models: thresholds and population regulation. Am. Nat. 121, 892–898. (doi:10.1086/284112)

3. Antonovics J, Iwasa Y, Hassell MP. 1995 A generalized model of parasitoid, venereal, and vector-based transmission processes. Am. Nat. 145, 661–675. (doi:10.1086/285761)

4. Begon M, Feore SM, Bown K, Chantrey J, Jones T, Bennett M. 1998 Population and transmission dynamics of cowpox in bank voles: testing fundamental assumptions. Ecol. Lett. 1, 82–86. (doi:10.1046/j.1461-0248.1998.00018.x)

5. Fenton A, Fairbairn JP, Norman R, Hudson PJ. 2002 Parasite transmission: reconciling theory and reality. J. Anim. Ecol. 71, 893–905. (doi:10.1046/j.1365-2656.2002.00656.x)

6. Ryder JJ, Webberley KM, Boots M, Knell RJ. 2005 Measuring the transmission dynamics of a sexually transmitted disease. Proc. Natl. Acad. Sci. U.S.A. 102, 15140–15143. (doi:10.1073/pnas.0505139102)

7. Smith MJ, Telfer S, Kallio ER, Burthe S, Cook AR, Lambin X, Begon M. 2009 Host– pathogen time series data in wildlife support a transmission function between density and frequency dependence. Proc. Natl. Acad. Sci. U.S.A. 106, 7905–7909. (doi:10.1073/pnas.0809145106)

8. Borremans B, Reijniers J, Hens N, Leirs H. 2017 The shape of the contact–density function matters when modelling parasite transmission in fluctuating populations. Roy. Soc. Open Sci. 4, 171308. (doi:10.1098/rsos.171308)

9. Antonovics J, Amoroso CR, Bruns E, Hood M. 2022 Host density shapes the relative contribution of vector-based and aerial transmission of a pathogenic fungus. Ecology 104, e3970. (doi:10.1002/ecy.3970)

10. Afshar A, Eaglesome MD. 1990 Viruses associated with bovine semen. Vet. Bull. 60, 93–109.

11. Roche BM, Alexander HM, Maltby AD. 1995 Dispersal and disease gradients of anthersmut infection of Silene alba at different stages. Ecology 76, 1863–1871. (doi:10.2307/1940719)

12. Bastos ADS, Bertschinger HJ, Cordel C, Vuuren CDJ van, Keet D, Bengis RG, Grobler DG, Thomson GR. 1999 Possibility of sexual transmission of foot-and-mouth disease from African buffalo to cattle. Vet. Rec. 145, 77–79. (doi:10.1136/vr.145.3.77)

13. Vitale F, Viviano E, Perna AM, Bonura F, Mazzola G, Ajello F, Romano N. 2000 Serological and virological evidence of non-sexual transmission of human herpesvirus type 8 (HHV8). Epidemiol. Infect. 125, 671–675. (doi:10.1017/s0950268800004726)

14. Mulligan CJ, Norris SJ, Lukehart SA. 2008 Molecular studies in Treponema pallidum evolution: toward clarity? PLoS Neglect. Trop. D. 2, e184. (doi:10.1371/journal.pntd.0000184)

15. Lawrence P, Danet N, Reynard O, Volchkova V, Volchkov V. 2017 Human transmission of Ebola virus. Curr. Opin. Virol. 22, 51–58. (doi:10.1016/j.coviro.2016.11.013)

16. Lockhart AB, Thrall PH, Antonovics J. 1996 Sexually transmitted diseases in animals: ecological and evolutionary implications. Biol. Rev. Camb. Philos. 71, 415–471. (doi:10.1111/j.1469-185X.1996.tb01281.x)

17. Alexander HM, Thrall PH, Antonovics J, Jarosz AM, Oudemans PV. 1996 Population dynamics and genetics of plant disease: a case study of anther-smut disease. Ecology 77, 990–996. (doi:10.2307/2265569)

18. Bruns EL, Antonovics J, Carasso V, Hood M. 2017 Transmission and temporal dynamics of anther-smut disease (*Microbotryum*) on alpine carnation (*Dianthus pavonius*). J. Ecol. 105, 1413–1424. (doi:10.1111/1365-2745.12751)

19. Ryder JJ, Miller MR, White A, Knell RJ, Boots M. 2007 Host-parasite population dynamics under combined frequency- and density-dependent transmission. Oikos 116, 2017–2026. (doi:10.1111/j.2007.0030-1299.15863.x)

20. Antonovics J, Boots M, Abbate J, Baker C, McFrederick Q, Panjeti V. 2011 Biology and evolution of sexual transmission. Ann. NY Acad. Sci. 1230, 12–24. (doi:10.1111/j.1749-6632.2011.06127.x)

21. Caldwell HD et al. 2003 Polymorphisms in *Chlamydia trachomatis* tryptophan synthase genes differentiate between genital and ocular isolates. J. Clin. Invest. 111, 1757–1769. (doi:10.1172/jci17993)

22. Faris R, Andersen SE, McCullough A, Gourronc F, Klingelhutz AJ, Weber MM. 2019 *Chlamydia trachomatis* serovars drive differential production of proinflammatory cytokines and chemokines depending on the type of cell infected. Front Cell Infect Microbiol 9, 399. (doi:10.3389/fcimb.2019.00399)

23. Gheit T. 2019 Mucosal and cutaneous human papillomavirus infections and cancer biology. Front. Oncol. 9, 355. (doi:10.3389/fonc.2019.00355)

24. Lukehart SA, Giacani L. 2014 When Is Syphilis Not Syphilis? Or Is It? Sex Transm Dis 41, 554–555. (doi:10.1097/OLQ.0000000000000179)

25. Thrall PH, Antonovics J, Wilson WG. 1998 Allocation to sexual versus nonsexual disease transmission. Am. Nat. 151, 29–45. (doi:10.1086/286100)

26. McLeod DV, Day T. 2019 Why is sterility virulence most common in sexually transmitted infections? Examining the role of epidemiology. Evolution 73, 872–882. (doi:10.1111/evo.13718)

27. Carlsson-Graner U, Thrall PH. 2006 The impact of host longevity on disease transmission: host-pathogen dynamics and the evolution of resistance. Evol. Ecol. Res. 8, 659–675.

28. Bruns E. 2019 Effects of host lifespan on the evolution of age-specific resistance: a case study of anther-smut disease on wild carnations. In Wildlife Disease Ecology: Linking Theory to Data and Application (eds A Fenton, D Tompkins, K Wilson), pp. 161–186. Cambridge: Cambridge University Press. (doi:10.1017/9781316479964.006)

29. Ross R. 1916 An application of the theory of probabilities to the study of a priori pathometry. Part I. Proc. R. Soc. Lond. A 92, 204–230. (doi:10.1098/rspa.1916.0007)

30. Ross R, Hudson HP. 1917 An application of the theory of probabilities to the study of a priori pathometry. Part II. Proc. R. Soc. Lond. A 93, 212–225. (doi:10.1098/rspa.1917.0014)

31. Kermack WO, McKendrick AG. 1997 A contribution to the mathematical theory of epidemics. Proc. R. Soc. Lond. A 115, 700–721. (doi:10.1098/rspa.1927.0118)

32. Anderson RM, May RM. 1978 Regulation and stability of host-parasite population interactions: I. regulatory processes. J. Anim. Ecol. 47, 219–247. (doi:10.2307/3933)

33. Bailey VA. 1931 The interaction between hosts and parasites. Q. J. Math. os-2, 68–77. (doi:10.1093/qmath/os-2.1.68)

34. Nicholson AJ, Bailey VA. 1935 The balance of animal populations. Part I. P. Zool. Soc. Lond. 105, 551–598. (doi:10.1111/j.1096-3642.1935.tb01680.x)

35. Soetaert K, Petzoldt T, Setzer RW. 2010 Solving differential equations in R: package deSolve. J. Stat. Softw. 33, 1–25. (doi:10.18637/jss.v033.i09)

36. D’Amico V, Elkinton JS, Dwyer G, Burand JP, Buonaccorsi JP. 1996 Virus transmission in gypsy moths is not a simple mass action process. Ecology 77, 201–206. (doi:10.2307/2265669)

37. Barlow ND. 2000 Non-linear transmission and simple models for bovine tuberculosis. J. Anim. Ecol. 69, 703–713. (doi:10.1046/j.1365-2656.2000.00428.x)

38. Reeson AF, Wilson K, Cory JS, Hankard P, Weeks JM, Goulson D, Hails RS. 2000 Effects of phenotypic plasticity on pathogen transmission in the field in a Lepidoptera-NPV system. Oecologia 124, 373–380. (doi:10.1007/s004420000397)

39. Hopkins SR, Fleming-Davies AE, Belden LK, Wojdak JM. 2020 Systematic review of modelling assumptions and empirical evidence: does parasite transmission increase nonlinearly with host density? Methods Ecol. Evol. 11, 476–486. (doi:10.1111/2041-210x.13361)

40. Kemler M, Denchev TT, Denchev CM, Begerow D, Piątek M, Lutz M. 2020 Host preference and sorus location correlate with parasite phylogeny in the smut fungal genus *Microbotryum* (Basidiomycota, Microbotryales). Mycol. Prog. 19, 481–493. (doi:10.1007/s11557-020-01571-x)

41. Thrall PH, Antonovics J. 1997 Polymorphism in sexual versus non-sexual disease transmission. Proc. R. Soc. Lond. B 264, 581–587. (doi:10.1098/rspb.1997.0083)

